# Loss of Flotillin-2 enhances trastuzumab emtansine internalization and cytotoxicity by relieving negative regulation of HER2 internalization in HER2-amplified cancers

**DOI:** 10.64898/2026.05.15.725439

**Authors:** David J. Wisniewski, Rachel K. Pritz, Jake Munch, Diya Desai, Tzu-Ting Huang, Sachin Kumar Deshmukh, Sharon Wu, Laurent Désaubry, George W. Sledge, Jung-Min Lee, Natalie Porat-Shliom, Stanley Lipkowitz

## Abstract

While Trastuzumab emtansine (T-DM1) and other HER2-targeting antibody-drug conjugates (ADCs) are used to treat cancer patients with HER2-amplified tumors, there is a need to improve the efficacy through the understanding of their mechanism of uptake into cells. Flotillin-2 (FLOT2) regulates the internalization of epidermal growth factor receptor (EGFR), leading us to investigate FLOT2 effects on HER2 internalization. Higher FLOT2 expression in nine HER2 amplified cell lines correlated with a higher T-DM1 IC50 *in vitro*, and breast cancer patients with high FLOT2 expression had worse survival when receiving either T-DXd (16.2 months (m) vs 18.3 m, p=0.04) or T-DM1 (38.0 m vs 41.3 m, p=0.1) in real-world Caris Life Sciences data. FLOT2 interacts with HER2 and positively regulates HER2 activation and downstream signaling, while FLOT2 knockdown reduces the viability of HER2 amplified cancer cells. FLOT2 knockdown results in increased HER2 internalization upon binding of T-DM1, mediated by ubiquitination by the Cbl ubiquitin ligases. We investigated the effects of various small molecules and discovered that zoledronic acid binds to FLOT2 and disrupts the HER2/FLOT2 interaction, which enhances T-DM1 internalization and cytotoxicity. In conclusion, FLOT2 regulates the internalization and cytotoxicity of T-DM1 mediated by Cbl-dependent ubiquitination of HER2. Zoledronic acid disrupts the HER2/FLOT2 interaction, therefore increasing the internalization and cytotoxicity of T-DM1, providing proof of principle that a small molecule inhibitor of the HER2/FLOT2 interaction can enhance the activity of the HER2-targeting ADC.

## Introduction

HER2 (ErbB2) is amplified in breast, gastric, uterine, biliary tract, colorectal, non-small cell, and bladder cancers (1). Considering the overexpression of this receptor and the reliance of the tumor on HER2 signaling, multiple HER2-targeted therapies have been developed clinically, including monoclonal antibodies, tyrosine kinase inhibitors, and antibody drug conjugates (ADCs) (1).

Flotillin-1 (FLOT1) and FLOT2 proteins are highly associated with lipid raft domains, and are described to regulate endocytosis, signal transduction, actin remodeling, and carcinogenesis (2–6). FLOT2 expression has been associated with poor prognosis and reduced survival in various cancers, including breast cancer and gastric cancer, both of which have HER2 amplified subtypes (7, 8). Our group previously described that FLOT2 regulates EGFR internalization and signaling, leading us to hypothesize that FLOT2 also regulates HER2 internalization and signaling (9). Indeed, it has been demonstrated that FLOT1 and FLOT2 interact with HER2, regulate HER2 expression, and are co-amplified with HER2 (7, 8, 10). Although FLOT2 and ErbB2 (the gene expressing HER2 protein) are located in the same region of the human chromosome, 17q11.2, their amplification is independent, as there are large regions of unamplified DNA between the two (11). FLOT1 and FLOT2 have also been described to reduce HER3 levels, and FLOT2 regulates AKT, MAPK, and FOXO signaling (12, 13). Interestingly, FLOT2 also regulated primary and metastatic growth of breast cancer cells without HER2 amplification *in vitro* and *in vivo* (11).

Despite the evidence that FLOT2 associates with and regulates HER2 expression, the detailed mechanism remains to be described, and the effects of FLOT2 on proliferation and growth has not been studied in HER2 amplified cancer. This study set out to determine the effects of FLOT2 on HER2 amplified cancer growth, the mechanism by which FLOT2 regulates HER2 internalization, and whether this mechanism is therapeutically exploitable. We focused on HER2 ADCs, studying trastuzumab emtansine (T-DM1) *in vitro*, and trastuzumab deruxtecan (T-DXd) using patient data. Our data shows that FLOT2 regulates HER2 and T-DM1 internalization mediated by the ubiquitin ligase Cbl. We also found that the small molecule zoledronic acid disrupts the HER2/FLOT2 interaction and potentiates the internalization and toxicity of T-DM1.

## Materials and Methods

### Materials

Fetal bovine serum (FBS) (631106) was obtained from Takara Bio. 4-20% precast polyacrylamide gels (5671094) were purchased from Bio-Rad. Immobilon PVDF transfer membrane (IPVH00010), mini protease inhibitor cocktail (11836153001), DMSO (D2650). Duolink In Situ Mounting Medium with DAPI (DUO82040), and propidium iodide (P4864) were purchased from Millipore-Sigma. PBS (21-031-CV) was purchased from Corning. Sodium orthovanadate (P0758S) was from Fisher Chemicals. NP40 Cell Lysis Buffer (FNN0021), Penicillin-Streptomycin (15140-122), DMEM (11965-092), 1M HEPES (15630-080), Sodium Pyruvate (11360-070), RPMI 1640 GLUTAMAX medium (61870036) and RPMI 1640 medium (11875-093) were obtained from ThermoFisher Scientific. Doxycycline Hyclate (198955) was purchased from MP Biomedicals. T-DM1 (ICH4014) was purchased from ichorbio. TAK-243 (S8341) was purchased from Selleck Chemicals. Protein A/G PLUS-Agarose (sc-2003) was purchased from Santa Cruz Biotechnology. Zoledronic Acid was obtained from Westminster Pharmaceuticals.

### Antibodies

Immunoblot antibodies: anti-phospho-HER2 Y1221/1222 (2243S), anti-phospho-HER2 Y1196 (6942S), anti-phospho-AKT S473 (9271S), anti-AKT (9272S), anti-phospho-MAPK T202/Y204 (9101), anti-phospho-S6 S235/236 (4856S), anti-S6 (2217S), anti-FLOT2 (3436S), anti-PHB1 (2426S) and anti-PHB2 (14085S) were purchased from Cell Signaling Technology. anti-HSC70 (sc-7298), anti-HA Tag (sc-7392), anti-Cbl (sc-393543), anti-Cbl-b (sc-8006), anti-Ubiquitin (sc-8017) and anti-ERK2 (D-2, sc-1647) antibodies were purchased from Santa Cruz Biotechnology. Anti mouse-IgGΚ BP-HRP (sc-516102) from Santa Cruz Biotechnology was used as the secondary antibody to develop immunoprecipitation bands for anti-FLAG and anti-FLOT2 because it does not interact with the IgG heavy chain. Goat Anti-Mouse IgG HRP conjugate (172-1011) and Goat Anti-Rabbit IgG HRP conjugate (1721019) were purchased from BioRad. Anti-HER2 (Rb-103-P1) was purchased from Neomarkers. anti-FLAG (F1804) was purchased from Millipore-Sigma. Immunoprecipitation antibodies: anti-FLAG (F1804) from Millipore-Sigma, anti-HER2 (sc-33684), anti-HA (sc-7392) and anti-FLOT2 (sc-28320) were purchased from Santa Cruz Biotechnology.

### Cell Lines

The HER2 amplified breast cancer cell lines SKBR3, HCC1954, BT474, HCC1419, HCC2218, UACC893 were obtained from ATCC. The HER2-amplified endometrial cancer cell lines ARK and ARK2 were kindly provided by Dr. Alessandro D. Santin, Yale University School of Medicine. All cell lines were maintained in RPMI 1640 medium with 10% FBS and 1% penicillin/streptomycin, and grown at 37°C, 5% CO_2_. HCC202 were obtained from ATCC and maintained in RPMI 1640 GLUTAMAX with 10% FBS, 1% penicillin/streptomycin, 1 mM Sodium Pyruvate and 25 mM HEPES. HeLa CRISPR control or FLOT2 knockout were generated as previously described (9) and were maintained in DMEM with 10% FBS and 1% penicillin/streptomycin, and grown at 37°C, 5% CO_2._ HEK293T were obtained from ATCC, maintained in DMEM with 10% FBS and 1% penicillin/streptomycin, and grown at 37°C, 5% CO_2._

### siRNA transfection

siNeg (4390844), siFLOT2 (s5285), siCbl (s2476), siCbl-b (s2479), siPHB1 (s10424) and siPHB2 (s22343) were all purchased from Thermo Fisher Scientific. Lipofectamine RNAiMAX (13778075) was purchased from Thermo Fisher Scientific, and transfections were performed according to manufacturer instructions. Of note, twice as much siFLOT2 as was recommended was used for each transfection.

### shRNA transduction

SMARTvector inducible Non-targeting Control mCMV/TurboGFP (shControl; VSC6570) and three separate SMARTvector Inducible shFLOT2 mCMV/TurboGFP sequences were purchased from Horizon Discovery/Dharmacon (V3SH7669-225045778; V3SH7669-226171705; V3SH7669-226615093) were purchased from Horizon Discovery/Dharmacon. Stable, selected pools for SKBR3, HCC202 and HCC1954 were transduced and selected according to manufacturer protocol. Expression of the shRNA was induced in the pools with doxycycline, cells were lysed and the lysates immunoblotted to determine the pool with optimal knockdown efficiency. The optimal knockdown pool was used for further testing. SKBR3 knockdown was most efficient with V3SH7669-226171705. HCC1954 knockdown was most efficient with V3SH7669-225045778. HCC202 knockdown was most efficient with V3SH7669-226615093. shControl promoter activity was confirmed by GFP visualization upon doxycycline treatment.

### Immunoblot and Immunoprecipitation

Lysates were collected, prepared and blotted as previously described (14). Densitometric analysis of band intensities for at least three independent experiments was calculated using ImageJ and data are presented as average ± SEM. Immunoprecipitation was performed as previously described (9).

### Cell Viability

Acridine Orange/Propidium Iodide staining was performed as previously described (9). Where mentioned, CellTiter-Glo 2.0 (Promega) was performed according to manufacturer instructions.

### Proximity Ligation Assay

Proximity ligation assay for FLOT2 and HER2 using probes conjugated with either anti-FLOT2 (F1680) from Millipore-Sigma or anti-HER2 (Rb-103-P1) from Neomarkers was performed as previously described (9).

### Caspase 3/7 Assay

Briefly, 5000 cells were plated and allowed to attach overnight and then treated in 1% FBS RPMI with the indicated conditions for the indicated time point. Following treatment, samples were incubated with Caspase 3/7 reagent (G8093) from Promega according to the manufacturer protocol.

### Dead Cell Assay

Propidium iodide (PI) staining, imaging and quantification was performed using the BioTek Cytation 1 Imaging Reader from Agilent (Santa Clara, CA) and Gen5 Image Prime 3.14 software (Agilent). Dead cell percentage was then calculated according to the manufacturer instructions. For PI, 1500-5000 cells were plated in a 96-well plate and treated the next day with experimental conditions in 1% FBS RPMI, along with PI (1:3000 dilution). Dead cell percentage was calculated by dividing dead cell number (PI stained) by brightfield cell count number and multiplying by 100 and normalized relative to the Day 0 dead cell percentage.

### Plasmid Transfection

HA-tag Ubiquitin plasmid was kindly provided by Dirk Bohmann. pCMV6 (PS10001) vector control and FLOT2 (RC220884) were purchased from OriGene. HER2 WT was a gift from Mien-Chie Hung (Addgene plasmid # 16257)(15). Lipofectamine 3000 (L3000008) was purchased from Thermo Fisher Scientific, and transfections were performed according to manufacturer instructions. Plasmids were transfected for a total of 72 hours, with a media change 24 hours post-transfection.

### T-DM1 uptake

CellTracker Blue CMAC (C2110) and pHrodo iFL Red Microscale Protein Labeling Kit (P36014) were purchased from Thermo Fisher Scientific. T-DM1 was conjugated to the pHrodo dye according to the manufacturer protocol. 100 ug of T-DM1 (1 mg/mL) was labeled with 3.3 uL of the dye. 25,000 cells were plated on Lab-Tek chambered coverglass (Thermo Fisher #155383) and allowed to attach overnight. The cells were then treated in experimental conditions. Then, the media was aspirated and cells were treated with the labeled T-DM1 (1 ug/mL) for 6.5 hours in serum-free RPMI. The media was then aspirated and the CellTracker Blue CMAC dye (1 µM) was added in serum-free media for 30 minutes. The media was then aspirated and fresh serum-free media without any additions was added, and images were acquired using Leica SP8 inverted confocal laser scanning microscope using 63×/1.4 objective. In ImageJ, individual cells were identified and manually selected using the CellTracker Blue CMAC and the cell area was saved with the ROI manager. The pHrodo fluorescence signal was thresholded, and the intensity in each cell was measured. For each cell, the fluorescence intensity was normalized to cell area and plotted relative to the control. Experiments were performed three times, and representative images are shown.

### Thermal Shift

2 million SKBR3 cells were plated and lysed the following day. Lysates were treated with PBS (control) or 300 µM zoledronic acid for 1 hour. Lysates were split into 8 separate Eppendorf tubes, and heated with a heating block at 40, 43, 46, 49, 52, 55, 58 and 61°C for three minutes each. The tubes were centrifuged at 17,000 RPM for 15 minutes, and the supernatant was saved. 2x Laemmli Sample Buffer (Bio-Rad #1610737) with β-mercaptoethanol was added to the lysate and boiled for 5 minutes and then immunoblotted.

### Caris Real-world data

Caris CODEai™ clinico-genomic database containing insurance claims data was used to calculate real-world overall survival (OS) from the start of T-DXd or T-DM1 treatment until last contact. Patients without contact/claims data for a period of at least 100 days were presumed deceased. Kaplan-Meier estimates were calculated for molecularly defined patient cohorts. Hazard ratios (HR) were determined by the Cox Proportional Hazards model, and p-values by the log-rank test. Significance was determined as p < 0.05.

### Statistical Analysis

Student’s *t*-test with 2 tailed comparisons assuming equal variance or one-way ANOVA were performed as indicated. p-values of ≤0.05 were considered significant. N.s indicates not statistically significant, * indicates p<0.05, ** indicates p<0.01, *** indicates p<0.001 and **** indicates p<0.0001.

## Results

### HER2 and FLOT2 interact; FLOT2 knockdown reduces viability and downstream signaling in HER2 amplified cancer cells

To determine whether FLOT2 is physically associated with HER2 in HER2-amplified cancer cells, we first examined their interaction using complementary approaches. Proximity ligation assay demonstrated a robust signal when HER2 and FLOT2 probes were present indicating that HER2 and FLOT2 are within close spatial proximity of each other, while no signal detected with either probe alone (Figure 1A). Immunoprecipitation of FLOT2 from SKBR3 cell lysates co-immunoprecipitates HER2, however this co-immunoprecipitation is lost when FLOT2 is knocked down, showing the specificity of the pull-down (Figure 1B). This was confirmed in HeLa cells, where HER2 immunoprecipitation pulls-down FLOT2, and this pull-down is lost when FLOT2 is knocked out (Figure 1C). Lastly, Flag-tagged FLOT2 only pulls-down HER2 when both HER2 and FLOT2 were co-transfected in HEK-293T cells (Figure 1D; lane 4).

**Figure 1.**
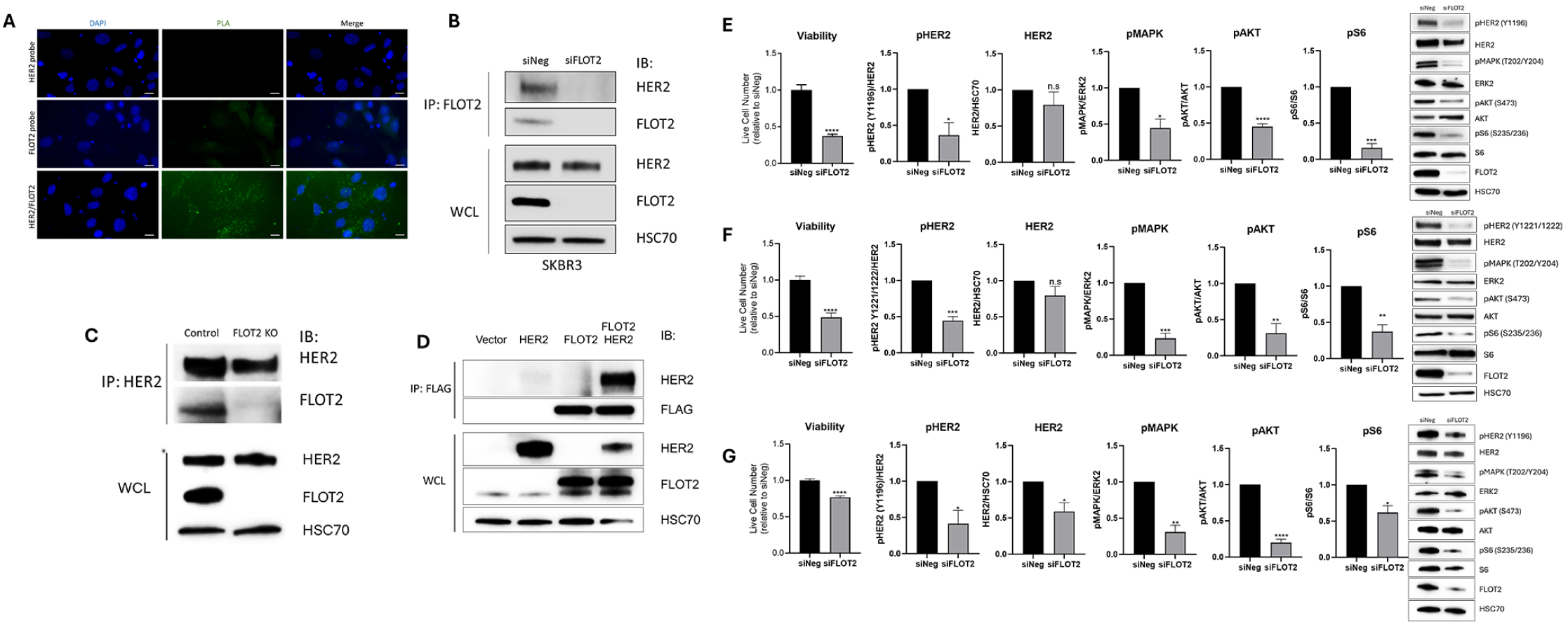
HER2 and FLOT2 interact; FLOT2 knockdown reduces viability and downstream signaling in HER2 amplified cancer cells. A) HCC1954 cells were fixed, permeabilized, blocked, and probed with PLA probes against HER2 and FLOT2 combined, or either probe alone (negative controls) and DAPI (blue). Green signal indicates interaction between HER2 and FLOT2. Scale bar is 10 µm. B) SKBR3 cells were transfected with siNeg or siFLOT2 for seven days prior to cell lysing and immunoprecipitation with anti-FLOT2, followed by immunoblotting for HER2 or FLOT2. Whole cell lysate was immunoblotted for HER2, FLOT2 or HSC70 (loading control). C) HeLa control or FLOT2 KO cells were lysed and immunoprecipitated with anti-HER2, followed by immunoblotting for HER2 and FLOT2. Whole cell lysate was immunoblotted for HER2, FLOT2 and HSC70 (loading control). D) HEK293T cells were transfected with empty vector, HER2, FLAG-tagged FLOT2 or HER2 and FLAG-tagged FLOT2 for three days, then lysed and immunoprecipitated with anti-FLAG and immunoblotted for HER2 and FLAG. Whole cell lysate was immunoblotted for HER2, FLOT2 and HSC70 (loading control). E) SKBR3 cells were transfected with siNeg or siFLOT2 for five days prior to cell counting or three days prior to lysing and immunoblot to probe for pHER2 (Y1196), HER2, pMAPK (T202/Y204), ERK2, pAKT (S473), AKT, pS6 (S235/236), S6, FLOT2, and HSC70 (loading control). Graphed data represent the average ±SEM of at least three independent experiments, and statistical analysis was performed by Student’s t-test. F) Same as in E, with HCC202 cells transfected for seven days prior to cell counting or three days prior to lysing, and lysates were instead probed for pHER2 (Y1221/1222). G) Same as in E, with HCC1954 cells transfected for seven days prior to cell counting and three days prior to lysing.

Previous studies in HER2 amplified breast cancer showed that FLOT2 regulated HER2 expression and AKT and MAPK signaling (7, 12). Our work confirmed that FLOT2 knockdown by either siRNA (Figure 1E-G) or doxycycline-induced shFLOT2 (Supplemental Figure S1) reduced the viability of three HER2 amplified breast cancer cell lines (SKBR3, HCC202 and HCC1954) as well as phosphorylated HER2, phosphorylated MAPK, phosphorylated AKT and phosphorylated S6. Of note, the knockdown of FLOT2 in HCC1954 by shFLOT2 was less dramatic, which may contribute to the less dramatic effect of knockdown on viability and downstream signaling (Supplemental Figure S1F). Doxycycline in shControl cells did not statistically significantly affect viability or HER2 activation or downstream signaling (Supplemental Figure S2). Interestingly, while pHER2 levels were reduced in all of the cells tested upon loss of FLOT2, the level of total HER2 was not consistently decreased in most of the cell lines by loss of FLOT2, suggesting that FLOT2 loss was predominantly causing loss of the activated HER2.

### FLOT2 knockdown increases TDM1 internalization and inhibits downstream signaling in HER2 amplified cells

We previously showed that FLOT2 regulates EGFR activation and internalization, (9). HER2 is a member of the EGFR family of receptors. Considering the interaction between HER2 and FLOT2, and the effect of FLOT2 on HER2 signaling, we hypothesized that FLOT2 regulates HER2 internalization. To evaluate T-DM1 internalization, we first conjugated T-DM1 to a pHrodo dye, which fluoresces in acidic cell compartments (e.g., endosomes and/or lysosomes), which signifies drug internalization. SKBR3 shFLOT2 cells treated with doxycycline to knock down FLOT2 had increased T-DM1 uptake, as visualized by an increased pHrodo TDM1 signal (Figure 2A). Green fluorescent protein (GFP) is downstream of the inducible promoter, indicating that the promoter is active upon doxycycline treatment. Importantly, doxycycline treatment in shControl cells did not significantly affect T-DM1 uptake, confirming that the increase in uptake in the shFLOT2 cells is due to FLOT2 knockdown, and not doxycycline alone (Figure 2B).

**Figure 2.**
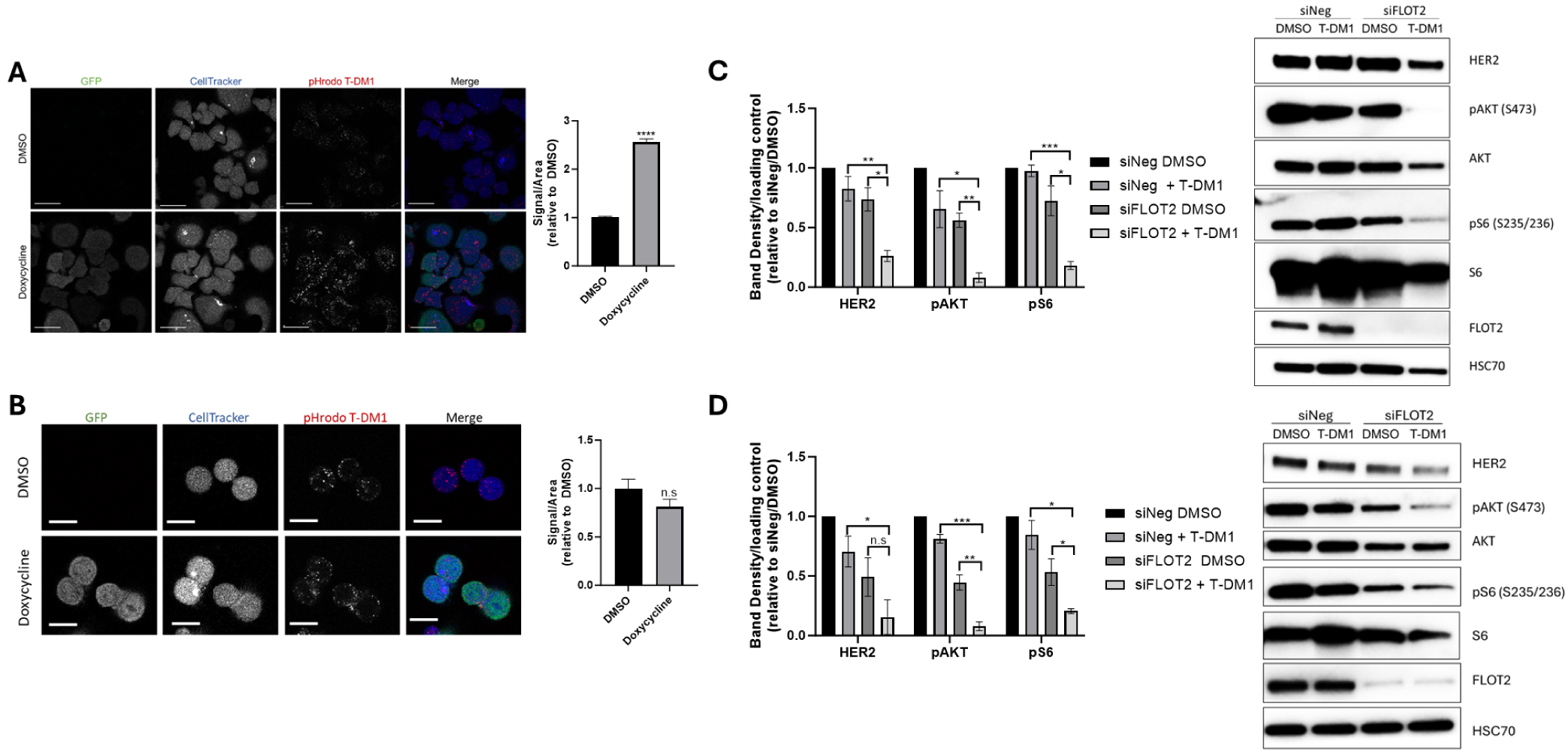
FLOT2 knockdown increases T-DM1 internalization and inhibits downstream signaling in HER2 amplified cells. A) SKBR3 shFLOT2 cells were treated +/- 500 ng/mL doxycycline for 72 hours, replated, then treated with pHrodo-T-DM1 (1 µg/mL; red) in serum-free RPMI for 7 hours. Cells were stained with CellTracker Blue CMAC dye (1 µM; Blue) in serum-free RPMI for 30 minutes prior to confocal imaging. GFP (green) expression is induced by doxycycline, indicating induction of the shFLOT2 promoter. Intensity of pHrodo signal per cell was quantified and divided by the area of each cell (right). Data represents the average ±SEM of at least three independent experiments, and statistical analysis was performed by Student’s t-test. Scale bar is 40 µm. B) Same as in A, with SKBR3 shControl cells. GFP (green) expression is induced by doxycycline, indicating induction of the shControl promoter. Scale bar is 20 µm. C) SKBR3 cells were transfected with siNeg or siFLOT2 for 48 hours and then treated with DMSO or 1 ng/mL T-DM1 for 24 hours in 1% FBS RPMI. Cells were lysed and immunoblotted for HER2, pAKT (S473), AKT, pS6 (S235/236), S6, FLOT2 and HSC70 (loading control). Band density was normalized to loading control, and then normalized to siNeg DMSO for each individual protein. Data represents the average ±SEM of at least three independent experiments, and statistical analysis was performed by Student’s t-test. D) Same as in C, with HCC202 cells using 10 µg/mL T-DM1.

We next examined the functional consequences of FLTOT2 depletion on HER2 signaling upon treatment with T-DM1. We treated two HER2 amplified breast cancer cells (SKBR3 and HCC202) with T-DM1 and FLOT2 knockdown and observed that total HER2, phosphorylated AKT and phosphorylated S6 were statistically significantly reduced compared to T-DM1 alone or FLOT2 knockdown alone (Figure 2C and D). In HCC202, total HER2 levels were lower with FLOT2 knockdown and T-DM1 treatment, however it was not statistically significant compared to FLOT2 knockdown alone (Figure 2D). Trastuzumab is known to increase HER2 activation potentially explaining the increase in total HER2 down regulation seen here compared to FLOT2 KD alone (16). Overall, FLOT2 knockdown dramatically increases TDM1 internalization, which in turn increases the ability of T-DM1 to inhibit HER2 signaling in HER2 amplified breast cancer cells.

### FLOT2 protein expression affects T-DM1 cytotoxicity in HER2 amplified cancer cells, while FLOT2 gene expression correlates with patient response to T-DM1 or T-DXd

We next hypothesized that the increase in T-DM1 internalization would result in increased drug cytotoxicity. We treated doxycycline-inducible shFLOT2 SKBR3 cells with doxycycline to knockdown FLOT2 and then treated with a T-DM1 dose curve to determine the IC50 of T-DM1 with FLOT2 knockdown (Figure 3A). Cells with less FLOT2 protein (gray) due to shFLOT2 induction had a 7.7 fold lower IC50 (1.5 ng/mL) compared to DMSO control cells (shFLOT2 induced 1.5 ng/ml vs control 11.53 ng/ml) as determined by cell viability (Figure 3A). This effect was not due to doxycycline cytotoxicity, as shControl cells exhibited a 1.6 fold increase in IC50 upon the induction of the shControl (shControl induced 57.66 ng/ml vs control 34.96 ng/ml). FLOT2 knockdown in HCC202 exhibited a 13 fold lower IC50 as compared to control (shFLOT2 induced 9.5 ng/ml vs control 125.2 ng/ml). The induced shControl HCC202 cells exhibited a much smaller 1.6 fold difference in IC50 (shControl induced 54.81 ng/ml vs control 71.29 ng/ml; Figure 3B). Similarly, FLOT2 knockdown in HCC1954 also exhibited a 11.1 fold lower IC50 as compared to control (shFLOT2 induced 4.6 ng/ml vs control 51.0 ng/ml), while shControl cells exhibited a much lesser 1.3 fold difference in IC50 (shControl induced 59.84 ng/ml vs control 78.1 ng/ml; Figure 3C). To determine whether this effect extends beyond breast cancer models, we examined whether this effect is observed in HER2 amplified uterine serous carcinoma cell lines ARK1 and ARK2. siRNA-mediated knockdown of FLOT2 reduced the IC50 of T-DM1 in ARK1by 2.2 fold compared to control siRNA (20.4 ng/ml in the siFLOT2 cells vs 45.0 ng/ml in siNEG), as well as in ARK2 (20.4 ng/mL in siFLOT2 cells compared to 40.6 ng/mL in siNEG cells) (Figure 3D). The viability comparison of siNeg vs siFLOT2 at multiple concentrations was statistically significant for both ARK1 and ARK2, even though the IC50 was not affected as much as the breast cancer cell lines. The difference in degree of sensitization in the HER2 amplified endometrial cancer cells may reflect the short term KD using siRNA vs the longer term KD using shRNA in the breast cancer cells. This effect is FLOT2 specific, as FLOT1 knockdown did not affect T-DM1 cytotoxicity in SKBR3 cells (Supplemental Figure S4).

**Figure 3.**
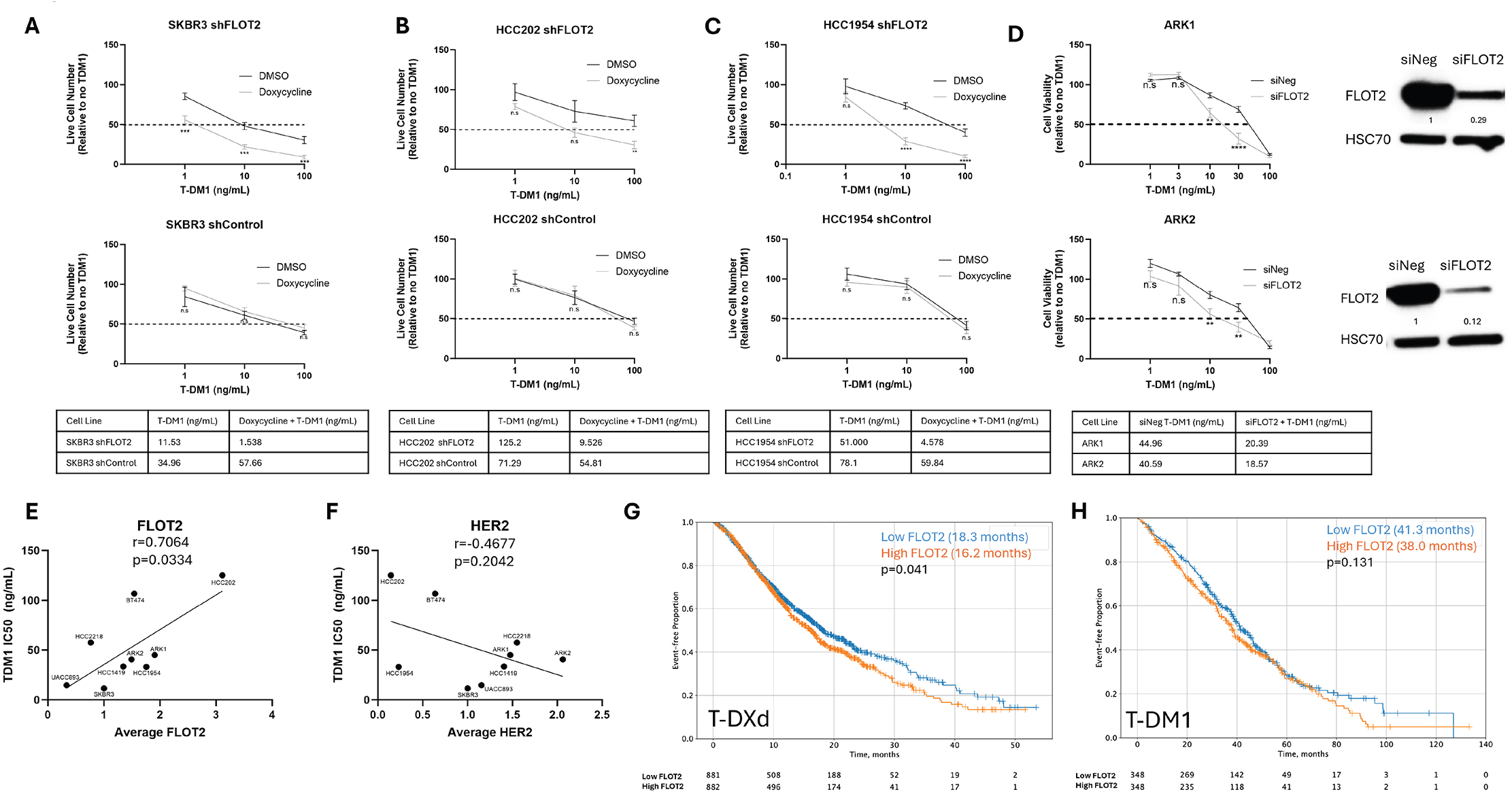
FLOT2 protein expression affects T-DM1 cytotoxicity in HER2 amplified cancer cells, while FLOT2 gene expression correlates with patient response to T-DM1 or T-DXD. A) SKBR3 shFLOT2 or shControl were pretreated with DMSO control (black) or 500 ng/mL doxycycline (gray) for 48 hours in 1% FBS RPMI, then treated with indicated T-DM1 concentration with continued doxycycline in 1% FBS RPMI for three days prior to cell counting. The table below the graphs indicates the calculated IC50 comparing DMSO + T-DM1 to doxycycline + T-DM1. Data represents the average ±SEM of at least three independent experiments, and statistical analysis was performed by Student’s t-test. B) Same as in A, with HCC202 shFLOT2 or shControl. C) Same as in A, with HCC1954 shFLOT2 or shControl. D) ARK1 (top) or ARK2 (bottom) cells were transfected with siNeg (black) or siFLOT2 (gray) for 48 hours prior to being treated with indicated T-DM1 concentration in 10% FBS RPMI for three days prior to cell viability reading (CellTiter-Glo 2.0). Data represents the average ±SEM of at least three independent experiments, and statistical analysis was performed by Student’s t-test. E) SKBR3, HCC202, HCC1954, BT474, HCC1419, HCC2218, UACC893, ARK1, and ARK2 cells were treated with 0-1000 ng/mL T-DM1 in 1% FBS RPMI for three days prior to cell counting. Average FLOT2 expression by immunoblot (relative to HSC70 loading control) and the calculated IC50 for each cell line was plotted. Protein expression and IC50 were calculated from at least three independent experiments. Line represents linear regression. F) Same as E, with average HER2 expression (relative to HSC70 loading control). G) Overall survival of patients in Caris dataset from T-DXd to last contact with high FLOT2 (orange; 16.22 months) vs low FLOT2 (blue; 18.26 months), p=0.041. H) Overall survival of patients in Caris dataset from T-DM1 to last contact with high FLOT2 (orange; 37.967 months) vs low FLOT2 (blue; 41.29 months), p=0.131.

Since FLOT2 knockdown enhances the effect of T-DM1 on HER2 amplified cancer cell viability, we investigated whether FLOT2 regulated the effect of T-DM1 on caspase activity, which is an indicator of apoptosis. FLOT2 knockdown by doxycycline in SKBR3, HCC202 and HCC1954 cells all statistically significantly increased caspase activity induced by T-DM1 compared to T-DM1 alone in the absence of FLOT2 knockdown (Supplemental Figure S5). Doxycycline treatment in shControl cells did not statistically significantly increase caspase activity in response to T-DM1 in SKBR3 (Supplemental Figure S5A, right panel) or HCC1954 (Supplemental Figure S5C, right panel). There was a statistically significant difference in HCC202 shControl comparing doxycycline with T-DM1 to T-DM1 alone (Supplemental Figure S5B, right panel), however the magnitude of the increase in the control cells compared to the shFLOT2 cells is dramatically different. In sum, loss of FLOT2 significantly increases T-DM1-mediated caspase activity in HER2 amplified breast cancer cells.

Higher FLOT2 expression in the HER2 amplified cell lines significantly correlated with a higher T-DM1 IC50 (r=0.7; p=0.03) (Figure 3E). In contrast, higher HER2 expression showed a nonsignificant correlation with lower T-DM1 IC50 in vitro (r=-0.5, p=0.2) (Figure 3F). Together, these results indicate that FLOT2 expression modulates cellular sensitivity to T-DM1

### Clinical correlation of FLOT2 expression with HER2-targeted ADC response

To explore the potential clinical relevance of the findings of the relationship between the efficacy of HER2 ADC’s and FLOT2 expression, we analyzed a large real world patient dataset treated with HER2-targeted ADCs from Caris Life Sciences (Caris). Patients with higher FLOT2 expression had a statistically significantly worse overall survival compared to low FLOT2 (16.2 months vs 18.3 months; p=0.04) in response to T-DXd, from onset of treatment to patient last contact (Figure 3G). Consistent with our in vitro data showing that high FLOT2 expression correlated with higher T-DM1 IC50, patients with higher FLOT2 expression had a borderline statistically significant worse overall survival compared to low FLOT2 (38.0 vs 41.3 months; p=0.1) in response to T-DM1 from onset of treatment to patient last contact (Figure 3H).

Given that FLOT2 and HER2 are often co-amplified (7, 8, 10), we examined survival outcomes with T-DXd or T-DM1 in samples with high or low HER2 expression levels. Patients with high HER2 had significantly better overall survival when compared to low HER2 (21.7 months vs. 13.4 months; p<0.00001) in response to T-DXd, from onset of treatment to patient last contact (Supplemental Figure S3A). Similarly, patients with high HER2 had significantly better overall survival when compared to low HER2 (46.5 months vs. 35.5 months; p<0.001) in response to T-DM1, from onset of treatment to patient last contact (Supplemental Figure S3B). Considering the co-amplification of FLOT2 and HER2, it is noteworthy that high HER2 has better outcomes when treated with T-DXd and T-DM1, but high FLOT2 has worse outcomes when treated with T-DXd and T-DM1.

We next investigated outcomes in high HER2/high FLOT2 compared to high HER2/low FLOT2 patients from onset of T-DXd treatment to patient last contact, since Caris had more patient data in response to T-DXd (Supplemental Figure S3C). Patients had significantly worse overall survival with high HER2/high FLOT2 compared to high HER2/low FLOT2 (18.5 months vs 25.6 months; p<0.001). Similarly, patients had statistically significantly worse overall survival with low HER2/high FLOT2 compared to low HER2/low FLOT2 (11.9 months vs 15.1months; p=0.002). Together, these data indicate that high FLOT2 negatively affects patient response to T-DXd independent of the HER2 levels.

Lastly, we analyzed whether FLOT1 expression had an effect on overall survival with either T-DXd or T-DM1 (Supplemental Figures S3E and S3F) from onset of treatment to patient last contact. FLOT1 expression did not have any effect on patient overall survival in response to T-DXd (low FLOT1= 17 months, high FLOT1= 17.1 months; p=0.7) or T-DM1 (low FLOT1= 38.9 months, high FLOT1= 40.6 months; p=0.7).

### FLOT2 knockdown increases ubiquitination of HER2 by T-DM1; ubiquitin-dependent HER2 internalizastion, and T-DM1 efficacy

Recent studies have reported that that small molecule tyrosine kinase inhibitors increased HER2 ubiquitination and T-DM1 internalization, and regulated HER2 degradation, although they did not confirm the requirment of ubiquitination for internalization (17, 18). Considering these studies, and our own work that indicated that FLOT2 decreased EGFR ubiquitination by Cbl, we sought to determine whether FLOT2 regulates the cellular effects of T-DM1 through modulation of HER2 ubiquitination (9). Knockdown of FLOT2 by doxycycline increased immunoprecipitation of HER2 by the anti-HA antibody targeting ubiquitin (Figure 4A, compare lane 4 to lane 3), indicating that loss of FLOT2 increases HER2 ubiquitination in response to T-DM1. To study whether the cytotoxic effects of T-DM1 were ubiquitin dependent, we treated SKBR3 and HCC1954 cells with TAK-243, a ubiquitin-activating enzyme inhibitor, and observed that inhibition of the first step of the ubiquitin pathway statistically significantly reduced the cytotoxicity of T-DM1 (Figure 4B). Of note, althoughTAK-243 alone exhibited cytotoxic effects (compare Bar 1 and Bar 3 in each panel), it abrogated the enhanced cytotoxicity observed with T-DM1 treatment (compare Bar 3 and 4 in each panel).

**Figure 4.**
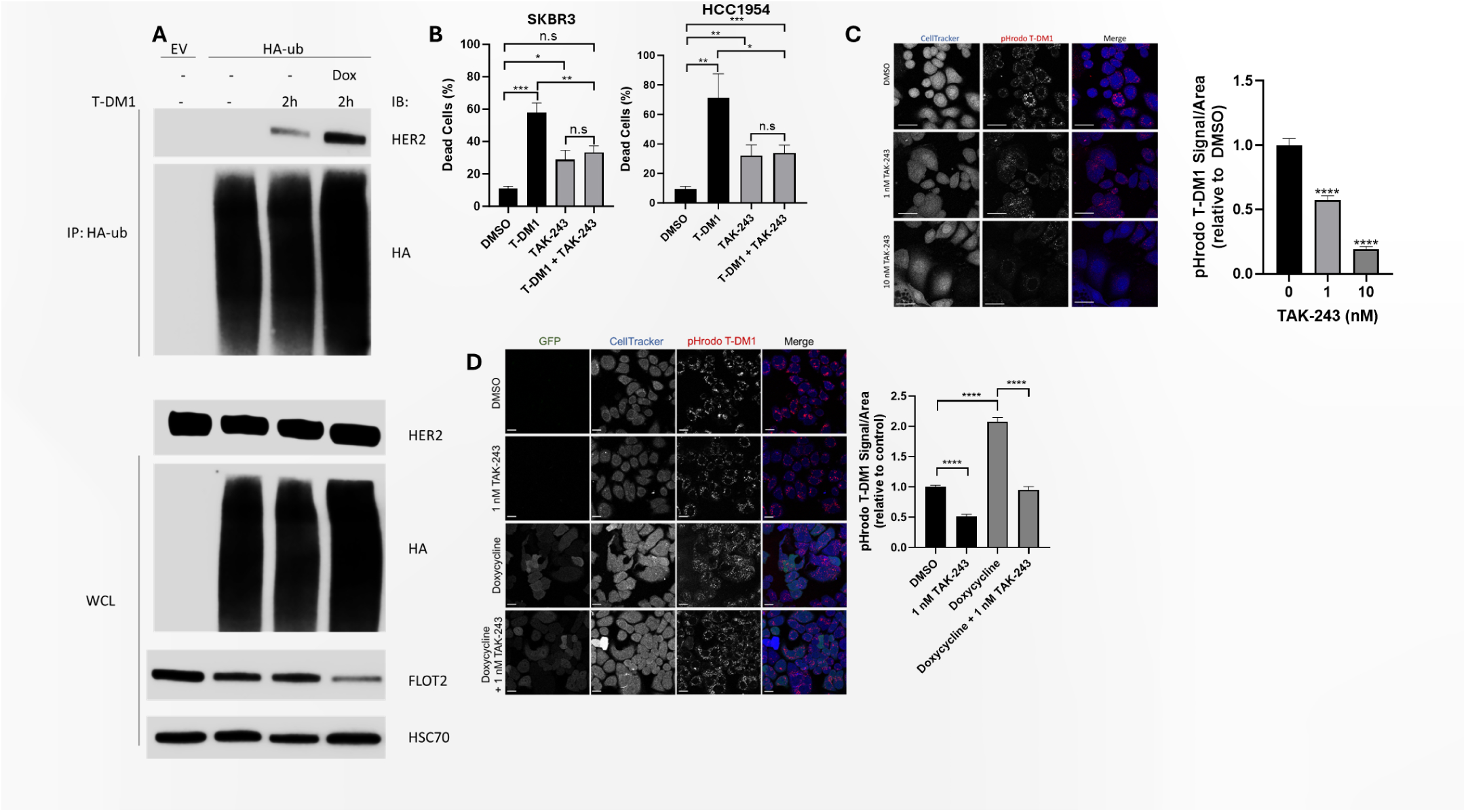
FLOT2 knockdown increases ubiquitination of HER2 by T-DM1; T-DM1 efficacy and internalization is ubiquitin-dependent. A) SKBR3 shFLOT2 cells were treated +/- 500 ng/mL doxycycline for a total of four days, and transfected with HA-tagged Ubiquitin for a total of three days. Cells were then treated with 10 ng/mL T-DM1 in 1% FBS RPMI for 2 hours, lysed, immunoprecipitated with anti-HA antibody, and immunoblotted for HER2 and HA. Whole cell lysate was immunoblotted for HER2, HA, FLOT2 and HSC70 (loading control). B) SKBR3 (left) or HCC1954 (right) were treated with TAK-243 (1 nM for SKBR3, 100 nM for HCC1954) and T-DM1 (10 ng/mL for SKBR3, 100 ng/mL for HCC1954) in 1% FBS RPMI for three days. Dead cell percentage was calculated using PI stain with the BioTek Cytation. Data represents the average ±SEM of at least three independent experiments, and statistical analysis was performed by Student’s t-test. C) SKBR3 cells were treated with DMSO (control), 1 nM or 10 nM TAK-243 for 24 hours in 1% FBS RPMI, and then treated with pHrodo-T-DM1 (1 µg/mL; red) for 7 hours in serum-free RPMI. Cells were stained with CellTracker Blue CMAC dye (1 µM; Blue) for 30 minutes in serum-free RPMI prior to confocal imaging. Intensity of pHrodo signal per cell was quantified and divided by the area of each cell (right). Data represents the average ±SEM of at least three independent experiments, and statistical analysis was performed by Student’s t-test. Scale bar is 40 µm. D) SKBR3 shFLOT2 were pretreated +/- 500 ng/mL doxycycline in complete RPMI for 48 hours, and then continued doxycycline +/- TAK-243 (1 nM) for 24 hours in 1% FBS RPMI. Cells were then treated with pHrodo-T-DM1 in serum-free RPMI (1 µg/mL; red) for 7 hours. Cells were stained with CellTracker Blue CMAC dye (1 µM; Blue) in serum-free RPMI for 30 minutes prior to confocal imaging. GFP (green) expression is induced by doxycycline, indicating induction of the shFLOT2 promoter. Intensity of pHrodo signal per cell was quantified and divided by the area of each cell (right). Data represents the average ±SEM of at least three independent experiments, and statistical analysis was performed by Student’s t-test. Scale bar is 20 µm.

To test whether the uptake of T-DM1 by cancer cells was also ubiquitin dependent, we treated SKBR3 cells with TAK-243 and subsequently treated with T-DM1-pHrodo (Figure 4C). Increasing concentrations of TAK-243 blocked T-DM1 internalization in a statistically significant manner as compared to control (Figure 4C, right). Using this same method in SKBR3 shFLOT2 cells, we observed that FLOT2 knockdown by doxycycline induction significantly increased T-DM1 uptake compared to control (Figure 4D, compare Bar 3 to Bar 1 in quantification), and this FLOT2-dependent effect was inhibited by TAK-243 treatment (Figure 4D, compare Bar 4 to Bar 3). Notably, TAK-243 also decreased the internalization of T-DM1 in control cells (Figure 4D, compare Bar 2 to Bar 1). Taken together, these data suggest that T-DM1 internalization is ubiquitin-dependent and FLOT2 negatively regulates ubiquitination of HER2 and T-DM1 uptake in the HER2 amplified SKBR3 cells.

### FLOT2 knockdown increases efficacy of T-DM1 in a Cbl and Cbl-b dependent manner

It has been previously established that the E3 ligase Cbl regulates HER2 ubiquitination (19, 20). Although Cbl may play a role in trastuzumab resistance, it has yet to be studied regarding T-DM1 or any other trastuzumab ADC internalization and efficacy in HER2 amplified cancer (21). We previously published that FLOT2 inhibits Cbl-mediated ubiquitination of EGFR in response to EGF (9). Based on these previous studies, we tested whether the increase in HER2 ubiquitination in response to T-DM1 treatment with FLOT2 knockdown was Cbl dependent. Consistent with the data shown in Figure 4 above, knockdown of FLOT2 resulted in increased immunoprecipitation of HER2 with the ubiquitin targeted HA antibody (Figure 5A, compare lane 5 to 3 in blot and bar 1 and 2 in graph). The increased ubiquitination of HER2 in response to FLOT2 knockdown cells treated with T-DM1 was inhibited by Cbl knockdown (Figure 5A, compare lane 10 to lane 5). This inhibition of T-DM1 induced HER2 ubiquitination by Cbl KD is significant when quantified (Figure 5A, graph, compare Bar 4 to Bar 2). In addition to the increased ubiquitination, the increased T-DM1 internalization with FLOT2 knockdown was inhibited by Cbl knockdown in SKBR3 cells (Figure 5B).

**Figure 5.**
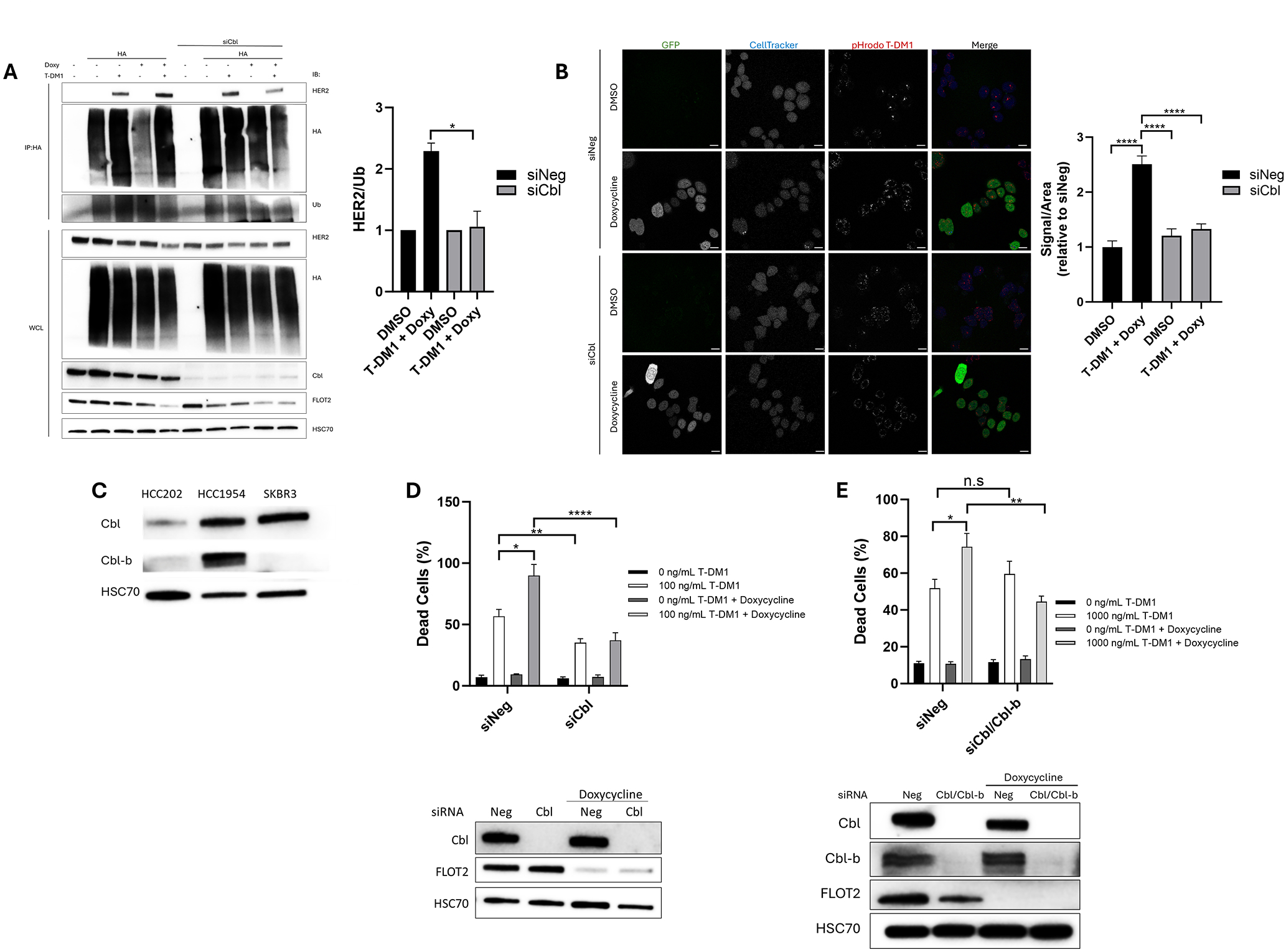
FLOT2 knockdown increases efficacy of T-DM1 in a Cbl and Cbl-b dependent manner. A) SKBR3 shFLOT2 cells were transfected with siRNA for Cbl (siCbl) or neg control (siNeg)and +/- 500 ng/mL doxycycline for 4 days total and transfected with HA-tagged ubiquitin for 3 days total. Cells were then treated with 10 ng/mL for 2 hours in 1% FBS RPMI, lysed, immunoprecipitated with anti-HA antibody, and immunoblotted for HER2, HA, and ubiquitin. Whole cell lysate was immunoblotted for HA, Cbl, FLOT2 and HSC70 (loading control). Immunoprecipitation band intensity was quantified and HER2 and ubiquitin and graphed as HER2/ubiquitin (right). siNeg with T-DM1 and doxycycline was normalized to siNeg with DMSO, and siCbl with T-DM1 and doxycycline was normalized to siCbl with DMSO. Data represents the average ±SEM of at least three independent experiments, and statistical analysis was performed by Student’s t-test. B) SKBR3 shFLOT2 cells were transfected with siNeg or siCbl +/- 500 ng/mL doxycycline for 48 hours and re-plated on chambered coverglass, then treated with pHrodo-T-DM1 (1 µg/mL; red) in serum-free RPMI for 7 hours. Cells were stained with CellTracker Blue CMAC dye (1 µM; Blue) in serum-free RPMI for 30 minutes prior to confocal imaging. GFP (green) expression is induced by doxycycline, indicating induction of the shFLOT2 promoter. Intensity of pHrodo signal per cell was quantified and divided by the area of each cell (right). Data represents the average ±SEM of at least three independent experiments, and statistical analysis was performed by one-way ANOVA. Scale bar is 20 µm. C) HCC202, HCC1954 and SKBR3 lysates were immunoblotted for Cbl, Cbl-b and HSC70 (loading control). D) The same SKBR3 shFLOT2 cells that were transfected with siNeg or siCbl and treated +/-doxycycline for 48 hours in B were then re-plated and treated +/- 500 ng/mL doxycycline +/- 100 ng/mL T-DM1 for an additional 72 hours and dead cell percentage was calculated using PI stain with the BioTek Cytation. Cells were lysed and immunoblotted (bottom) for Cbl, FLOT2 and HSC70 (loading control) to confirm knockdown. Data represents the average ±SEM of at least three independent experiments, and statistical analysis was performed by Student’s t-test. Dead cell percentage was normalized to the baseline percentage recorded at time of TDM1 treatment. E) Same as in D, with HCC1954 shFLOT2 cells. SiCbl was co-transfected with si-Cbl-b and cells were treated with 1000 ng/mL T-DM1, and lysates were also immunoblotted for cbl-b.

We next tested if the knockdown of Cbl proteinsabrogated the increased toxicity of T-DM1 seen upon FLOT2 loss in the HER2 amplified SKBR3 and HCC1954 cell lines. Of note, SKBR3 cells express Cbl but not Cbl-b protein, whereas HCC1954 express both Cbl and Cbl-b, therefore in SKBR3 we only knocked down Cbl, but did a dual knockdown of Cbl and Cbl-b in HCC1954 (Figure 5C). In SKBR3 cells, knockdown of Cbl abrogated the increased cytotoxicity of shFLOT2 combined with T-DM1 in a statistically significant manner (Figure 5D, top, compare Bar 8 to Bar 6)). Similarly, in HCC1954, knockdown of Cbl/Cbl-b abrogated the increased cytotoxicity of shFLOT2 combined with T-DM1 in a statistically significant manner (Figure 5E, top). FLOT2, Cbl and Cbl-b knockdown were confirmed by western blot (Figure 5D and E, bottom panels). Taken together, the internalization and cytotoxicity of T-DM1 is enhanced by FLOT2 knockdown, and this effect is through Cbl-mediated ubiquitination of HER2.

### Disruption of the HER2 and FLOT2 interaction by the small molecule zoledronic acid increases T-DM1 mediated toxicity

As a proof of principle, we investigated whether disrupting the HER2 and FLOT2 interaction using a small molecule could phenocopy the loss of FLOT2 on T-DM1 toxicity. Zoledronic acid has been reported to bind to PHB2, which shares a PHB domain with FLOT2 (22). The binding of zoledronic acid to FLOT2, PHB1 or PHB2 was investigated by thermal shift assay (Figure 6A). The melting temperature of both PHB2 and FLOT2 increased upon zoledronic acid incubation, consistent with protein stabilization by the drug. The melting temperature of PHB1 was unchanged, indicating no binding of zoledronic acid to PHB1. As FLOT1 did not affect T-DM1 cytotoxicity, we did not test the effects of zoledronic acid on FLOT1 stability (Supplemental Figure S4).

**Figure 6.**
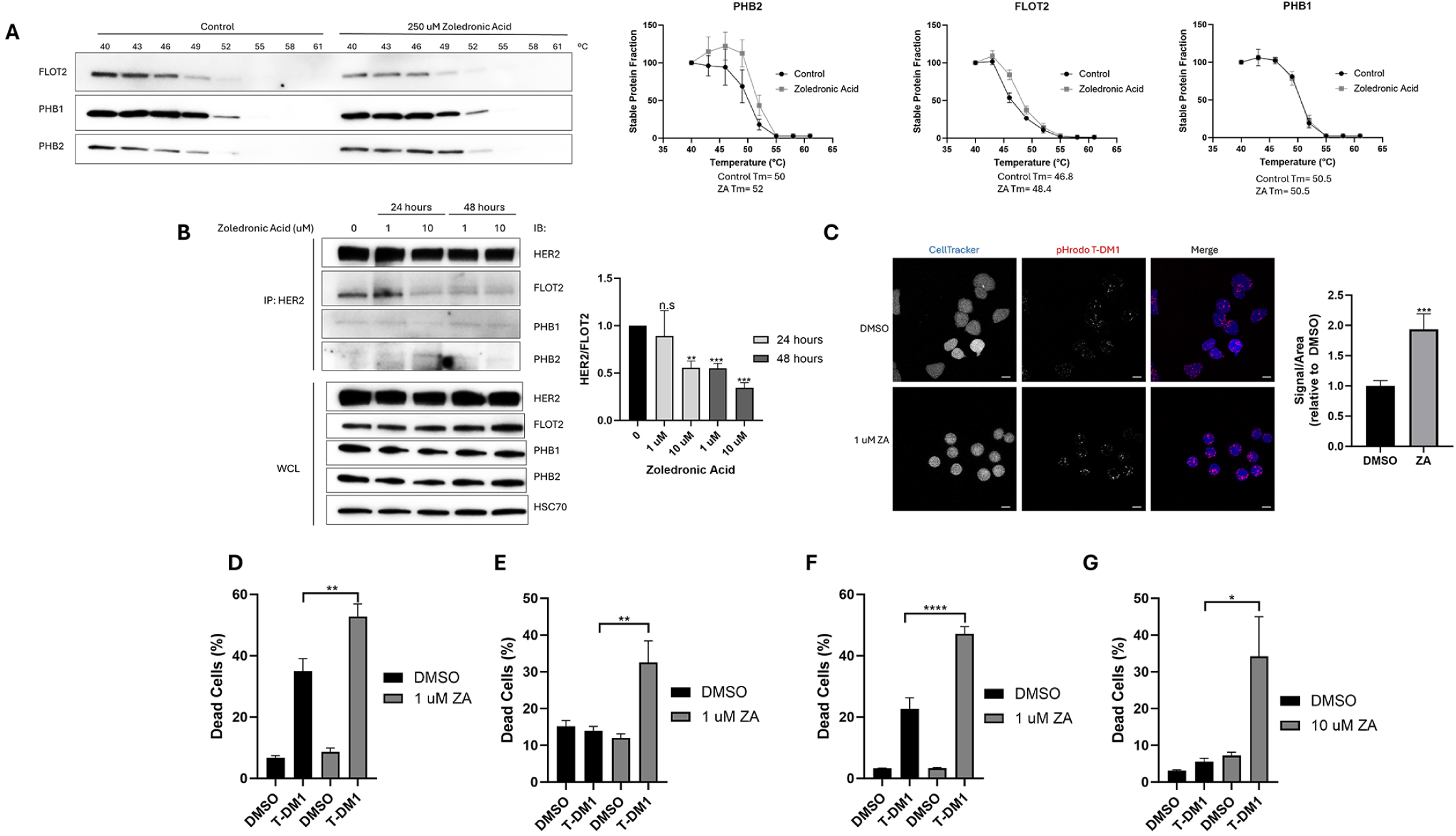
Zoledronic Acid binds to FLOT2, disrupting HER2 and FLOT2 interaction and increases T-DM1 cytotoxicity. A) SKBR3 cells were lysed, lysates treated with DMSO or 250 µM zoledronic acid (ZA) for an hour, and then heated at the indicated temperatures for 3 minutes each. Lysates were then centrifuged, and supernatant was immunoblotted for FLOT2, PHB1, and PHB2 (left). Quantification of bands and calculation of melting temperatures are depicted on the right. Quantification of bands represent the average ±SEM of three independent experiments. B) SKBR3 cells were treated with 0, 1 or 10 µM zoledronic acid for 24 or 48 hours, lysed, immunoprecipitated by anti-HER2, and immunoblotted for HER2, FLOT2, PHB1 or PHB2. Whole cell lysate was immunoblotted for HER2, FLOT2, PHB1, PHB2 and HSC70 (loading control). Quantification of immunoprecipitation bands from three independent experiments for FLOT2 relative to HER2 are depicted on the right, with the average ±SEM. Statistical analysis was performed by Student’s t-test compared to 0 µM zoledronic acid. C) SKBR3 cells were treated with DMSO (control) or 1 µM Zoledronic acid (ZA) for 72 hours then treated with pHrodo-T-DM1 (1 µg/mL; red) in serum-free RPMI for 7 hours. Cells were stained with CellTracker Blue CMAC dye (1 µM; Blue) in serum-free RPMI for 30 minutes prior to confocal imaging. Intensity of pHrodo signal per cell was quantified and divided by the area of each cell (right). Data represents the average ±SEM of at least three independent experiments, and statistical analysis was performed by Student’s t-test. Scale bar is 20 µm. D) SKBR3 cells were treated with the indicated combinations of DMSO (control), 1 ng/mL T-DM1 and 1 µM zoledronic acid (ZA) for 48 hours in 1% FBS RPMI. Dead cell percentage was calculated using PI stain with the BioTek Cytation. Data represents the average ±SEM of three independent experiments, and statistical analysis was performed by Student’s t-test. E) Same as D, with ARK1 cells in 10% FBS RPMI. F) Same as E, with ARK2 cells and 10 µM zoledronic acid (ZA). G) Same as D, with HCC1954 cells with 24 hour treatment.

We next examined whether the binding of zoledronic acid to FLOT2 would disrupt the interaction of FLOT2 with HER2 and hence would phenocopy FLOT2 knockdown. Indeed, zoledronic acid disrupted the co-immunoprecipitation of HER2 and FLOT2 in a statistically significant manner (Figure 6B; blot and graph). Zoledronic acid did not affect HER2 co-immunoprecipitation with either PHB1 or PHB2, as their co-immunoprecipitation was minimally detectable (Figure 6B, blot). This indicates that PHB1 and PHB2 minimally interact with HER2, and the binding of zoledronic acid to PHB2 is not affecting this HER2/FLOT2 interaction. Zoledronic acid treatment increased internalization of T-DM1, as the T-DM1-pHrodo signal was increased in the presence of zoledronic acid in a statistically significant manner (Figure 6C). Zoledronic acid treatment led to an increase in cytotoxicity of T-DM1, as treatment of SKBR3, ARK1, ARK2 and HCC1954 (Figure 6D-G, respectively) with T-DM1 and zoledronic acid all exhibited significant increases in cytotoxicity when compared to T-DM1 alone. Zoledronic acid alone did not exhibit cytotoxicity.

To further confirm that zoledronic acid’s effect on T-DM1 is FLOT2 dependent, and not a PHB1 or PHB2 effect, we knocked down either PHB1 or PHB2. PHB1 or PHB2 knockdown did not significantly increase T-DM1 cytotoxicity in SKBR3 cells (Supplemental Figure S6A). In fact, at 100 ng/mL T-DM1, PHB1 or PHB2 knockdown significantly reduced T-DM1 cytotoxicity as compared to siNeg control. Further, PHB1 or PHB2 knockdown did not affect HER2 and FLOT2 co-immunoprecipitation (Supplemental Figure S6B). Taken together, this indicates that zoledronic acid enhances T-DM1 cytotoxicity and internalization by binding to FLOT2 and disrupting the interaction between HER2 and FLOT2. Although zoledronic acid also binds to PHB2, the observed effects of zoledronic acid on FLOT2 and T-DM1 is independent of PHB1 or PHB2. Together, these data indicate that a small molecule can disrupt the FLOT2/HER2 interaction and potentiate the internalization and efficacy of a HER2 directed ADC.

## Discussion

The goals of this study were to elucidate the role of FLOT2 on HER2 signaling and internalization, it role in T-DM1 internalization and cytotoxicity, as well as the mechanism by which this is mediated. Further, we set out to determine if there are any compounds which may exploit this mechanism, resulting in improved T-DM1 internalization and cytotoxicity. We confirmed that HER2 and FLOT2 interact in HER2 amplified cancer cell lines (Figure 1 A-D), and FLOT2 protein expression positively regulates HER2 signaling and viability (Figure 1 E-G and Supplemental Figure S1). Higher FLOT2 expression reduces T-DM1 internalization, downstream signaling effects, cytotoxicity, and caspase activity (Figures 2, 3 and Supplemental Figure S5). High FLOT2 expression correlates with a higher T-DM1 IC50 *in vitro,* and breast cancer patients with high FLOT2 expression had worse overall survival in response to T-DM1 and T-DXd (Figure 3 G-H, Supplemental Figure S3C-D). FLOT2 reduces HER2 ubiquitination upon T-DM1 treatment, which results in reduced T-DM1 internalization (Figure 4). The increased T-DM1 internalization and cytotoxicity upon FLOT2 knockdown is dependent on the Cbl and Cblb ubiquitin ligases (Figure 5). Lastly, we demonstrated that zoledronic acid binds to FLOT2, disrupts the interaction between FLOT2 and HER2, resulting in increased T-DM1 internalization and cytotoxicity in HER2 amplified cancer (Figure 6).

The mechanism by which FLOT2 regulates HER2 is different than the mechanism previously published by our group regarding FLOT2 regulating EGFR (9). Although HER2 and EGFR are in the same family, their basal activity and conformations differ (23). EGFR typically exists as an inactive monomer, and upon ligand binding it dimerizes and becomes active, resulting in increased downstream signaling (24). The HER2 extracellular domain is structurally similar to the active EGFR extracellular domain and amplified HER2 exists basally as a catalytically active dimer (23). Therefore, we postulate that FLOT2 binds to the inactive EGFR monomer at the membrane, and upon FLOT2 knockdown there is increased ability of the EGFR to dimerize at the membrane, resulting in increased activation, ubiquitination, and internalization upon activation. Since HER2 is already activated and dimerized basally, FLOT2 binding to HER2 at the membrane does not prevent activity. Loss of FLOT2, however, leads to increased Cbl-mediated ubiquitination and internalization of the HER2 receptor.

Several studies have identified HSP90 as a regulator of HER2 and trastuzumab or T-DM1 internalization (7, 25–29). It has been demonstrated that HSP90 complexes with FLOT2 as well (7). These previous studies led us to test whether the HSP90 inhibitor geldanamycin could synergize with T-DM1 in a FLOT2-dependent manner. However, our own experimentation with multiple geldanamycin concentrations combined with multiple T-DM1 concentrations did not elicit any synergistic effects, and we therefore sought out alternative mechanisms to explain the role of FLOT2 on T-DM1 internalization and cytotoxicity.

The patient data indicates that high FLOT2 expression is associated with worse overall survival in response to T-DM1 and T-DXd (Figure 3 G-H, Supplemental Figure S3C-D). This study provides evidence that FLOT2 expression may stratify patients’ ability to respond to HER2 targeted ADCs.

This is not the first study to examine improving T-DM1 efficacy, as multiple studies have used small molecule tyrosine kinase inhibitors to increase T-DM1 internalization and efficacy (17, 18, 30). These studies, however, have differing conclusions whether they increase or decrease HER2 ubiquitination upon T-DM1 treatment, and did not use knockdown or inhibitors to determine whether the mechanism by which these inhibitors increase T-DM1 internalization is ubiquitin dependent or independent. Our study provides evidence HER2 internalization is E1-dependent and Cbl-dependent and that FLOT2 inhibits ubiquitination and internalization of HER2 (Figures 4 and 5). Future studies should fully determine the mechanisms by which these tyrosine kinase inhibitors regulate T-DM1 internalization, which may provide evidence for other pathways which can be targeted to further enhance the efficacy of T-DM1 and other HER2 target ADCs.

This study identified zoledronic acid as a compound which binds to FLOT2, disrupts the FLOT2/HER2 interaction, and increases the toxicity of the T-DM1 – providing proof of principle that a small molecule can phenocopy the loss of FLOT2 to improve HER2 directed ADC activity. Two small studies have tested adding zoledronic acid to neoadjuvant therapy for HER2 positive breast cancers with contradicting conclusions about benefit of the addition of zoledronic acid (31–33). However, these studies did not evaluate zoledronic acid in the context of HER2-directed ADCs. While the concentration of zoledronic acid that was effective at disrupting the FLOT2/HER2 complex and increasing the internalization and toxicity of T-DM1(e.g., 1 μM) is within the range of the Cmax of zoledronic acid in the blood (∼1 μM), the clearance of zoledronic acid from blood is very rapid with concentrations decreasing to <1% of the Cmax by 24 hours (34). In contrast, the half-life of ADCs such as T-DM1 and T-Dxd are typically 3-6 days (35, 36). Zoledronic acid incorporates into bone and is typically given to patients at 4-12 week intervals for bone metastases and at 6 month intervals in the adjuvant setting to prevent bone recurrences and bone mineral loss (37, 38). Thus, zoledronic acid would not be optimal for enhancing the efficacy of HER2-directed ADCs. Therefore, we propose that future studies expand the search for compounds which bind FLOT2 and disrupt its interaction with HER2, including potential analogs of zoledronic acid which may be more specific to disrupting the FLOT2/HER2 interaction and have better pharmacokinetic properties.

Overall, this study provides evidence that FLOT2 negatively regulates the ubiquitin dependent internalization and cytotoxicity of T-DM1. Further, zoledronic acid binds to FLOT2, disrupting the interaction between FLOT2 and HER2, leading to an increase in internalization and cytotoxicity of T-DM1. Taken together with patient data that low FLOT2 in breast cancer patients correlates with improved response to T-DM1 and T-DXd, compounds which disrupt FLOT2 and HER2 binding should be pursued for potential combination with HER2-targeted ADC’s.

## Acknowledgments

We thank all members of the Lipkowitz lab in the Women’s Malignancies Branch, Center for Cancer Research, National Cancer Institute for their support and review of this manuscript. This research was supported by the Intramural Research Program of the National Cancer Institute (ZIA BC 010977) of the National Institutes of Health (NIH). The contributions of the NIH authors are considered Works of the United States Government. The findings and conclusions presented in this paper are those of the authors and do not necessarily reflect the views of the NIH or the U.S. Department of Health and Human Services.

## Data Availability Statement

The Caris deidentified sequencing data are owned by Caris Life Sciences and cannot be publicly shared due to the data usage agreement in place. These data will be made available to researchers for replication and verification purposes through the Caris letter of intent process, which are generally fulfilled within 6 months. For more information on how to access this data, please contact Joanne Xiu at. All other data generated in this study are included in this published article and its supplementary information.

## Supplemental Figures

**S1. Inducible shFLOT2 knockdown reduces viability and downstream signaling in HER2 amplified cancer cells.** A) SKBR3 shFLOT2 cells were treated +/- 500 ng/mL doxycycline for seven days prior to cell counting and five days prior to cell lysis. shFLOT2 lysates were probed for pHER2 (Y1196), HER2, pMAPK (T202/Y204), ERK2, pAKT (S473), AKT, pS6 (S235/236), S6, FLOT2, and HSC70 (loading control). Graphed data represent the average ±SEM of at least three independent experiments, and statistical analysis was performed by Student’s t-test. B) Same as A, with HCC202 shFLOT2 cells, and lysates were instead probed for pHER2 (Y1221/1222). Cells were treated with doxycycline for seven days prior to cell counting and cell lysis. C) Same as in A, with HCC1954 shFLOT2 cells. Cells were treated with doxycycline for seven days prior to cell counting and cell lysis.

**S2. Doxycycline does not affect cell viability or HER2 signaling in shControl cells.** A) SKBR3 shControl cells were treated +/- 500 ng/mL doxycycline for seven days prior to cell counting and five days prior to cell lysis. shControl lysates were probed for pHER2 (Y1196), HER2, pMAPK (T202/Y204), ERK2, pAKT (S473), AKT, pS6 (S235/236), S6, FLOT2, and HSC70 (loading control). Data represents the average ±SEM of at least three independent experiments, and statistical analysis was performed by Student’s t-test. B) Same as A, with HCC202 shControl cells, and lysates were instead probed for pHER2 (Y1221/1222). Cells were treated with doxycycline for seven days prior to cell counting and cell lysis. C) Same as in A, with HCC1954 shControl cells. Cells were treated with doxycycline for seven days prior to cell counting and cell lysis.

**S3. Overall survival in response to T-DXd or T-DM1 in Caris data.** A) Overall survival of patients in Caris dataset from T-DXd to last contact with high HER2 (orange; 21.747 months) vs low HER2 (blue; 13.357 months), p<0.00001. B) Overall survival of patients in Caris dataset from T-DM1 to last contact with high HER2 (orange; 46.521 months) vs low HER2 (blue; 35.499 months), p<0.001. C) Overall survival of patients in Caris dataset from T-DXd to last contact with high HER2/high FLOT2 (blue; 18.457 months) vs high HER2/low FLOT2 (orange; 25.629 months), p<0.001. D) Overall survival of patients in Caris dataset from T-DXd to last contact with low HER2/low FLOT2 (blue; 15.134 months) vs low HER2/high FLOT2 (orange; 11.943 months), p=0.002. E) Overall survival of patients in Caris dataset from T-DXd to last contact with high FLOT1 (orange; 17.141 months) vs low FLOT1 (blue; 17.042 months), p=0.732. F) Overall survival of patients in Caris dataset from T-DM1 to last contact with high FLOT1 (orange; 40.631 months) vs low FLOT1 (blue; 38.921 months), p=0.682.

**S4. FLOT1 knockdown does not affect T-DM1 cytotoxicity in SKBR3 cells.** SKBR3 cells were transfected with siNeg or siFLOT1 for 48 hours, replated, and then treated with 0, 1, 10 or 100 ng/mL T-DM1 in 1% FBS RPMI for 72 hours. Dead cell percentage was calculated using PI stain with the BioTek Cytation. Data represents the average ±SEM of at least three independent experiments, and statistical analysis was performed by Student’s t-test, comparing individual TDM1 concentration of siFLOT1 to siNeg.

**S5. FLOT2 knockdown increases caspase activity in response to T-DM1.** A) SKBR3 shFLOT2 (left) or shControl (right) were treated +/- 500 ng/mL doxycycline for 48 hours and then treated with 10 ng/mL T-DM1 as indicated for 24 hours. Cells were then incubated with Caspase 3/7 reagent, luminescence was recorded, and values were normalized to DMSO control. Data represents the average ±SEM of at least three independent experiments, and statistical analysis was performed by Student’s t-test. B) Same as in A, with HCC202 shFLOT2 or shControl. C) Same as in A, with HCC1954 shFLOT2 or shControl, and 100 ng/mL T-DM1.

**S6. PHB1 or PHB2 knockdown does not affect HER2/FLOT2 co-immunoprecipitation or T-DM1 cytotoxicity.** A) SKBR3 cells were transfected with siNeg, siPHB1 or siPHB2 for 72 hours, replated, and treated with 0, 1, 10 or 100 ng/mL T-DM1 for 48 hours. Dead cell percentage was calculated using PI stain with the BioTek Cytation. Viable cell percentage was calculated by subtracting dead cell percentage from 100. These percentages were normalized to 0 ng/mL TDM1 for each transfection condition. Data represents the average ±SEM of at least three independent experiments, and statistical analysis was performed by Student’s t-test, comparing different T-DM1 concentrations in either siPHB1 or siPHB2 to siNeg T-DM1 concentrations. B) Transfected SKBR3 cells from A were lysed, immunoprecipitated for anti-HER2, and immunoblotted for HER2 and FLOT2. Whole cell lysate was immunoblotted for HER2, FLOT2, PHB1, PHB2 and HSC70 (loading control). Immunoprecipitation was quantified for HER2/FLOT2, and normalized to siNeg (right). Data represents the average ±SEM of at least three independent experiments, and statistical analysis was performed by Student’s t-test.

