## Supplementary figures and images for "Loss of Flotillin-2 enhances trastuzumab emtansine internalization and cytotoxicity by relieving negative regulation of HER2 internalization in HER2-amplified cancers"

### Supplementary Figure 1

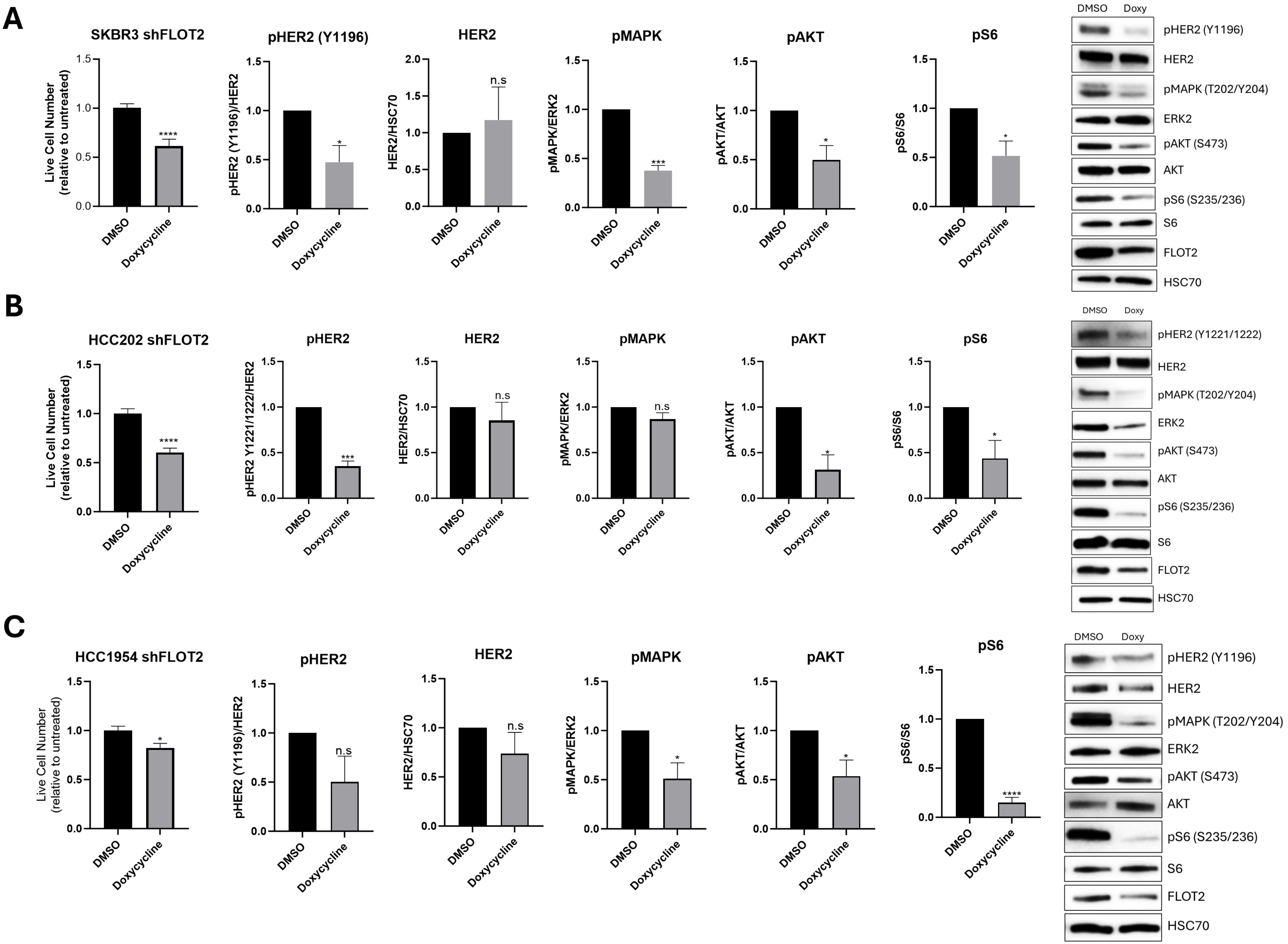

### Supplementary Figure 2

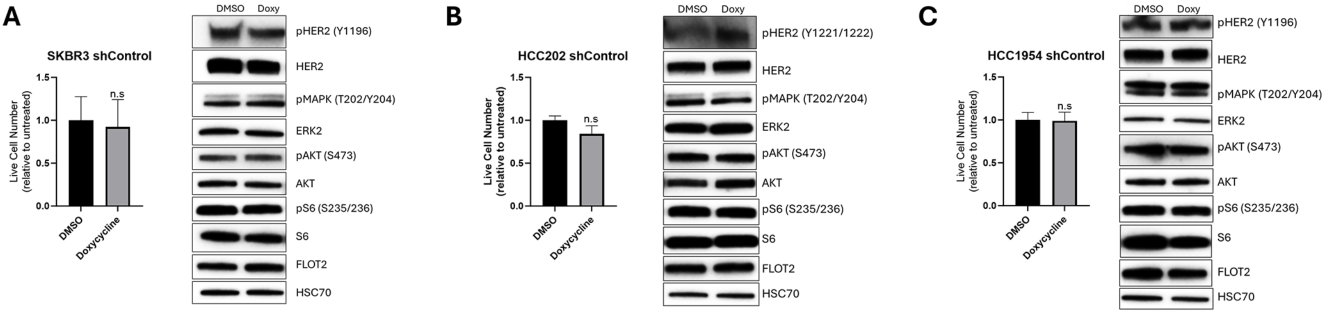

### Supplementary Figure 3

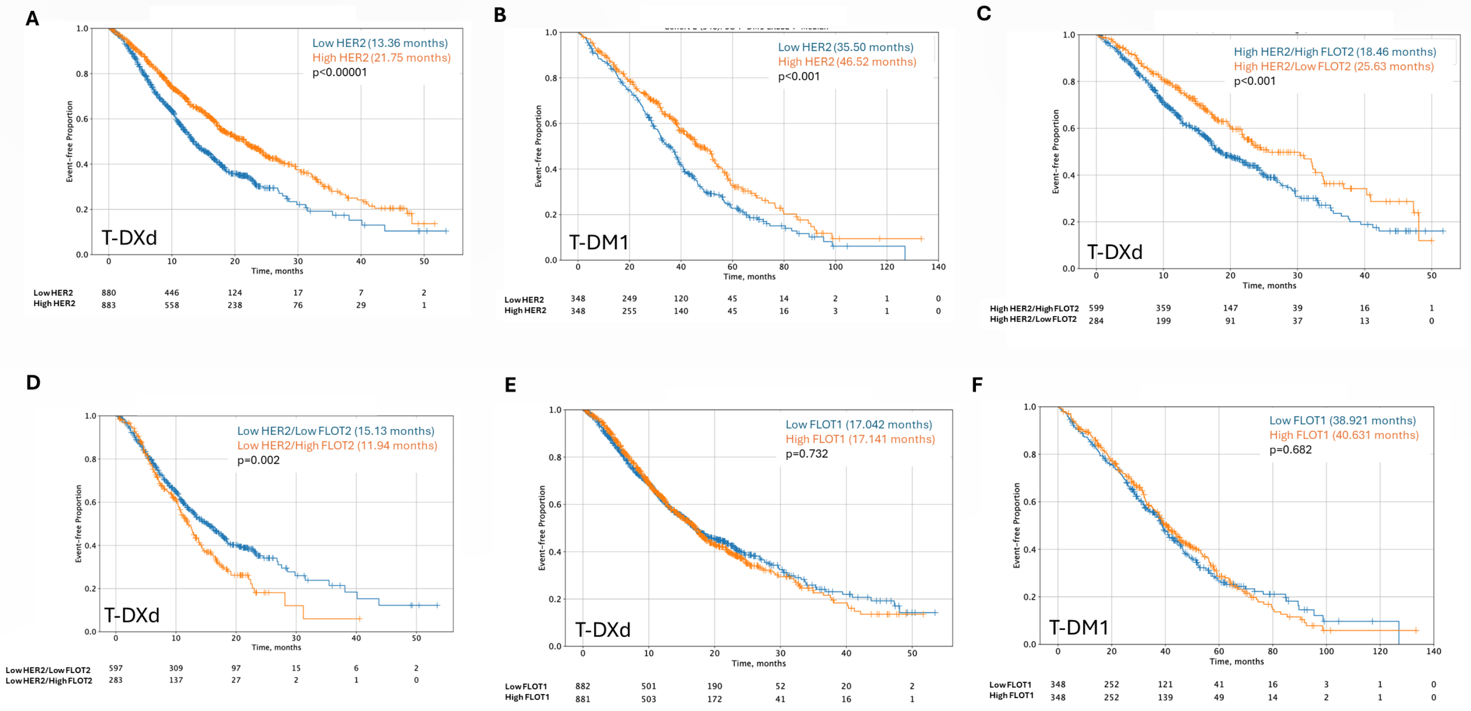

### Supplementary Figure 4

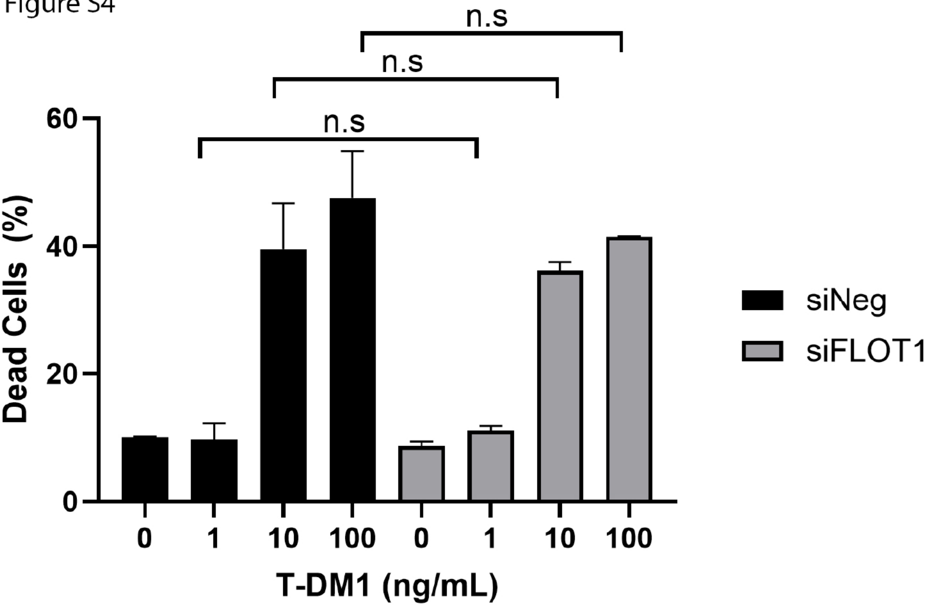

### Supplementary Figure 5

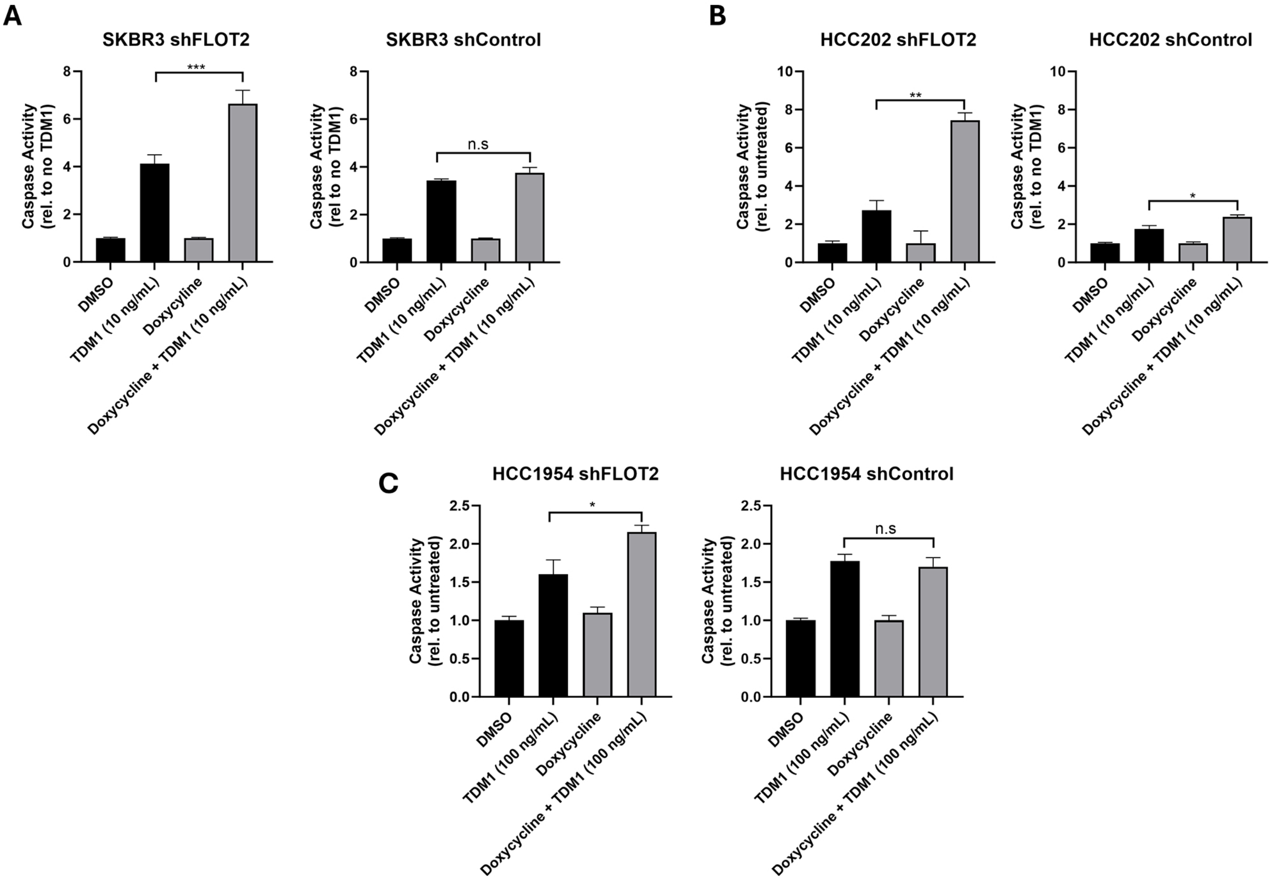

### Supplementary Figure 5

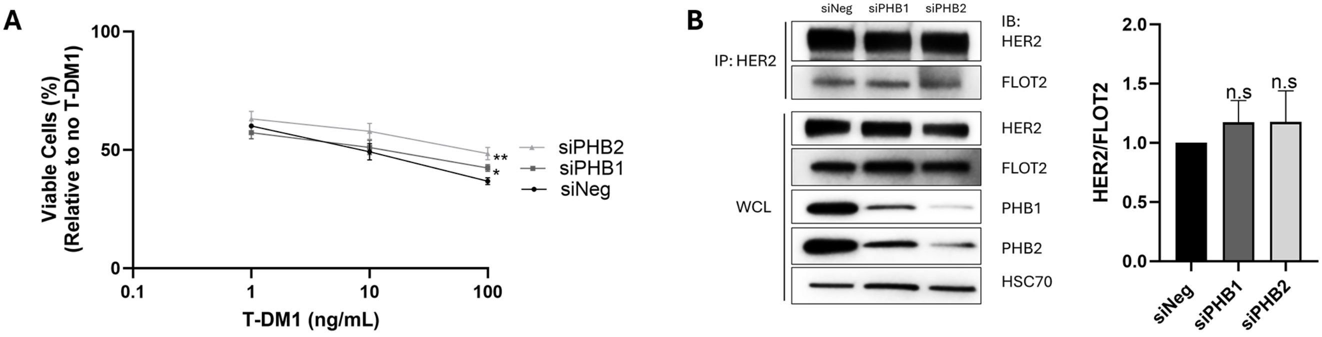
